# Applying a Modified Metabarcoding Approach for the Sequencing of Macrofungal Specimens from Fungarium Collections

**DOI:** 10.1101/2021.12.22.473928

**Authors:** C. Gary Olds, Jessie W. Berta-Thompson, Justin J. Loucks, Richard A. Levy, Andrew W. Wilson

## Abstract

- *Premise:* Fungaria are a largely untapped source for understanding fungal biodiversity. The effort and cost in producing DNA barcode sequence data for large numbers of fungal specimens can be prohibitive. This study applies a modified metabarcoding approach that provides a labor and cost-effective solution for sequencing the fungal DNA barcode from hundreds of specimens at once.
- *Methods:* A two-step PCR approach uses nested barcoded primers to nrITS2 sequence data. We applied this to 766 macrofungal specimens that represent a broad taxonomic sampling of the Dikarya, of which 382 *Lactarius* specimens are used to identify molecular operational taxonomic units (MOTUs) through a phylogenetic approach. Scripts in Python and R were used to organize sequence data and execute packages CutAdapt and DADA2 were used for primer removal and assessing sequence quality. Sequences were compared to NCBI and UNITE databases and Sanger-produced sequences.
- *Results:* Specimen taxonomic identities from nrITS2 sequence data are >90% accurate across all specimens sampled. Phylogenetic analysis of *Lactarius* sequences identified 20 MOTUs.
- *Discussion:* The results demonstrate the capacity of these methods to produce nrITS2 sequences from large numbers of fungarium specimens. This provides an opportunity to more effectively use fungarium collections in advancing fungal diversity identification and documentation.

## INTRODUCTION

Only 8% of the estimated 2.2 to 3.8 million fungal species worldwide have been documented (Hawksworth and Lücking, 2017). Our ability to address the gap between known and unknown fungal diversity will require new approaches in methodology and technology. Fungaria (herbaria of fungi) are an essential resource that preserve and provide the specimens necessary for investigating and documenting fungal diversity (Andrew et al., 2018). Many fungarium collections consist of macrofungi which are fungi that produce macroscopic reproductive structures (e.g. mushrooms, puffballs, club fungi, coral fungi, cup fungi, truffles). These fungi largely represent diverse taxonomic groups from the subkingdom Dikarya (phyla Ascomycota and Basidiomycota). The specimens from these collections provide opportunities for research on the taxonomy, systematic diversity, evolution, and ecology of fungi in both narrow and broad scopes (Andrew et al., 2018). However, fungaria remain a largely untapped resource for exploring questions in fungal biology, in large part because so many specimens have incomplete or suspicious identifications. Using DNA sequence data is a logical step towards identifying undescribed fungal specimens. However, there are several challenges with generating DNA sequence data from large numbers of specimens, which can be summed up in three parts: scale, sequencing effectiveness, and cost.

The challenge of scale is exemplified by the number of unidentified specimens in fungaria (for clarification we use “unidentified” to indicate “not identified to species”). The large number of unidentified fungal specimens is a major impediment for examining questions in fungal population genetics, ecology, and distributions. The Mycology Collections Portal (MyCoPortal) database for North American fungaria has records for 343,993 fungal specimens (Mycoportal.org, 2021). Of these, 77,897 (23%) are unidentified. The Global Biodiversity Information Facility (GBIF) database aggregates records of fungal specimens globally, containing over 7.5 million records (Gbif.org, 2021). Of these, slightly less than 3 million have the specific epithets, suggesting that around 60% of fungal specimens in global collections are unidentified.

The rather large number of unidentified macrofungal specimens historically comes from the reliance on morphology in fungal systematics. For fungal diversity studies, DNA barcoding has become as a necessary first step in species recognition (Lücking et al., 2020). The DNA barcode for fungi is the nuclear ribosomal internal transcribed spacer (nrITS) region (Schoch et al., 2012), which was selected because of the high PCR amplification success rate, coupled with the high interspecific but low intraspecific variation. Sequences of nrITS from many fungal specimens have been shown to be effective at recognizing useful molecular operational taxonomic units (MOTUs) in multiple taxonomic groups (Begerow et al., 2010; Osmundson et al., 2013). These features helped contribute toward making the nrITS the most common fungal sequence on GenBank (Lücking et al., 2020). Despite this, the vast majority of accepted fungal species do not have a representative nrITS sequence on Genbank (Hawksworth and Lücking, 2017). Compound this with the potentially large number of misidentified specimens in fungaria and the potential for identifying new species from macrofungal specimens becomes too significant to ignore. The ability of mycologists to efficiently produce DNA barcode data from hundreds to thousands of fungal specimens is a problem for which modern high-throughput sequencing technologies may offer solutions. However, these solutions have not yet been developed.

The Sanger sequencing approach to DNA barcoding fungal specimens has been used for over 3 decades (White et al., 1990), and many systematic and taxonomic studies currently rely on its effectiveness and practicality (Noffsinger and Cripps, 2021; Vera et al., 2021). Despite Sanger sequencing’s long productive history, there are some significant limitations to its effectiveness. One of these is the inability to sequence degraded DNA from older specimens. Additionally, translating the sequences into usable nrITS data can be complicated by irregular insertions and deletions of nucleotides among ribosomal copies resulting in frame shifts that are often interpreted as noise in the electropherograms. The issue of scale when producing nrITS data from large numbers of unidentified fungarium specimens quickly runs up against the barrier of cost using the Sanger method. For example, we are charged approximately $4.00 USD per sequence when we outsource Sanger sequencing. Forward and reverse sequence reads are generally produced for redundancy and to span the sequence length from both directions, making the cost of one nrITS contiguous sequence (or contig) per specimen approximately $8.00 USD. Scaling this cost to sequence thousands of unidentified specimens in fungaria quickly makes it prohibitive to perform comprehensive studies in fungal diversity.

High-throughput sequencing (e.g., pyrosequencing, next, second, and third generation sequencing) can address the challenges of scale, effectiveness, and cost when generating DNA barcode sequence data from fungal specimens. These sequencing technologies accomplish this by using different chemistries and data recording methods. Also, the sheer number of reads produced can be used for validating the sequence data. Metabarcoding studies use this technology to generate a portion of the fungal DNA barcode to document communities from soil and tissue samples (Nilsson et al., 2019). Given that a single study can produce sequence data representing hundreds to thousands of species, the method has potential for use on fungarium collections. Several versions of these methods typically produce short reads. Metabarcoding studies that use these short read sequencing methods need to focus on one of two regions in the nrITS, the ITS1 or ITS2. The utility of these markers for DNA barcoding can vary among taxa, but the overall differences in phylogenetic signal between the two are considered negligible and similar results can be obtained from either marker (Bazzicalupo et al., 2013; Blaalid et al., 2013). Between the two, the ITS2 region is slightly more favored for DNA barcoding given the number of reference sequences available (Nilsson et al., 2008; Lücking et al., 2020), and complications in targeting ITS1 in some taxa (Lindahl et al., 2013). In terms of cost, approaches like Illumina MiSeq Nano provide 800,000 read pairs for a little over $1000 USD, plus approximately $500 for library quantification, cleaning, and additional PCR through the sequencing facility (source: University of Idaho’s IIDS Genomics and Bioinformatics Resources Core). Spread this across 1000 fungarium specimens and one will theoretically average 800 reads/specimen at a sequencing cost between $1-$2/specimen (Figure 1).

**Figure 1.**
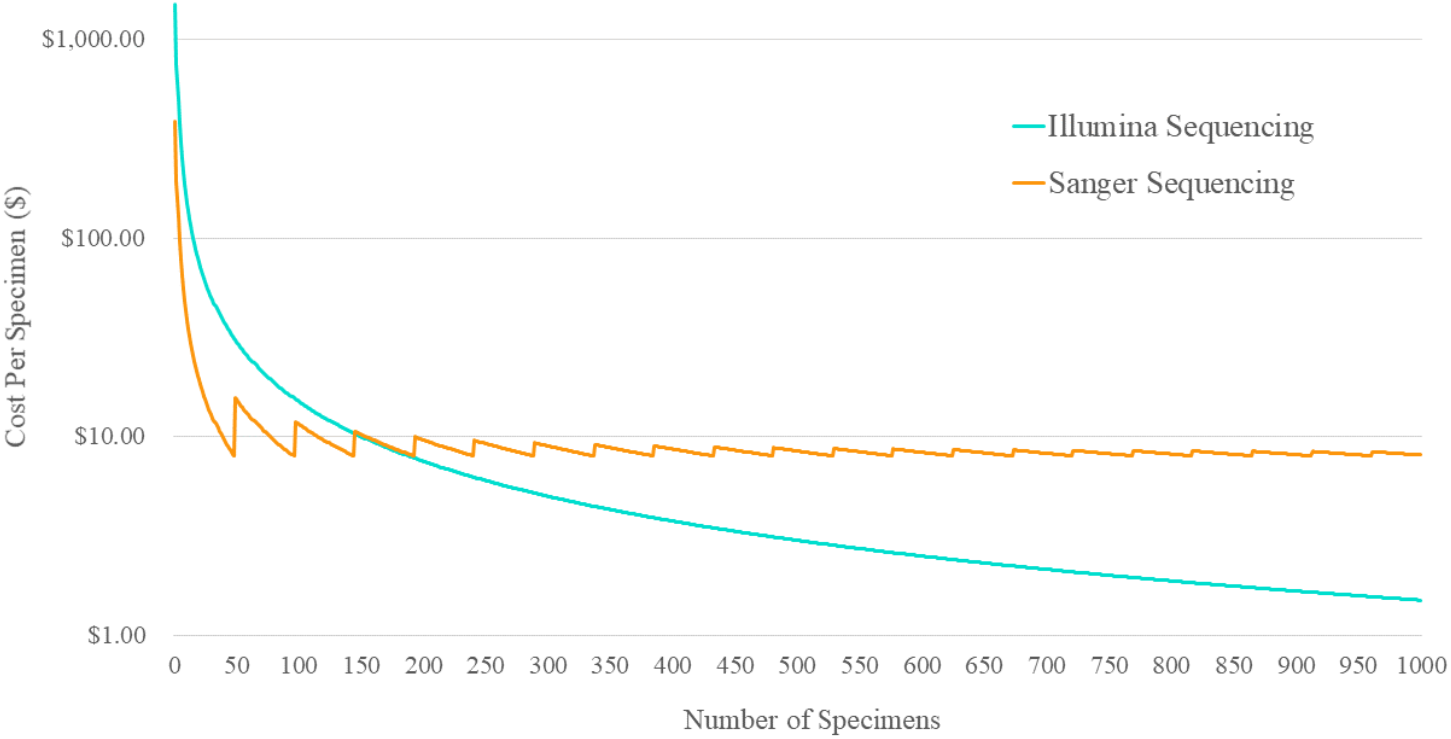
Cost comparison between Illumina and Sanger Sequencing for sequencing DNA barcodes from fungal specimens. Sanger sequencing costs are based on a rate of $4 per sequence. Illumina sequencing cost is based on a baseline cost of $1,500 for Illumina MiSeq Nano (500 cycles) sequencing and additional library prep.

Novel approaches of high-throughput sequencing have been developed for studying natural history collections (e.g., Folk et al., 2021). Currently, there is no high-throughput sequencing approach designed for the DNA barcoding of fungarium collections. In this project we have developed and tested a modified metabarcoding approach for sequencing the nrITS2 region in fungal specimens (Figure 2). We assess the utility of this approach across the subkingdom Dikarya, its effectiveness on older specimens, and its capacity to recognize MOTUs.

**Figure 2.**
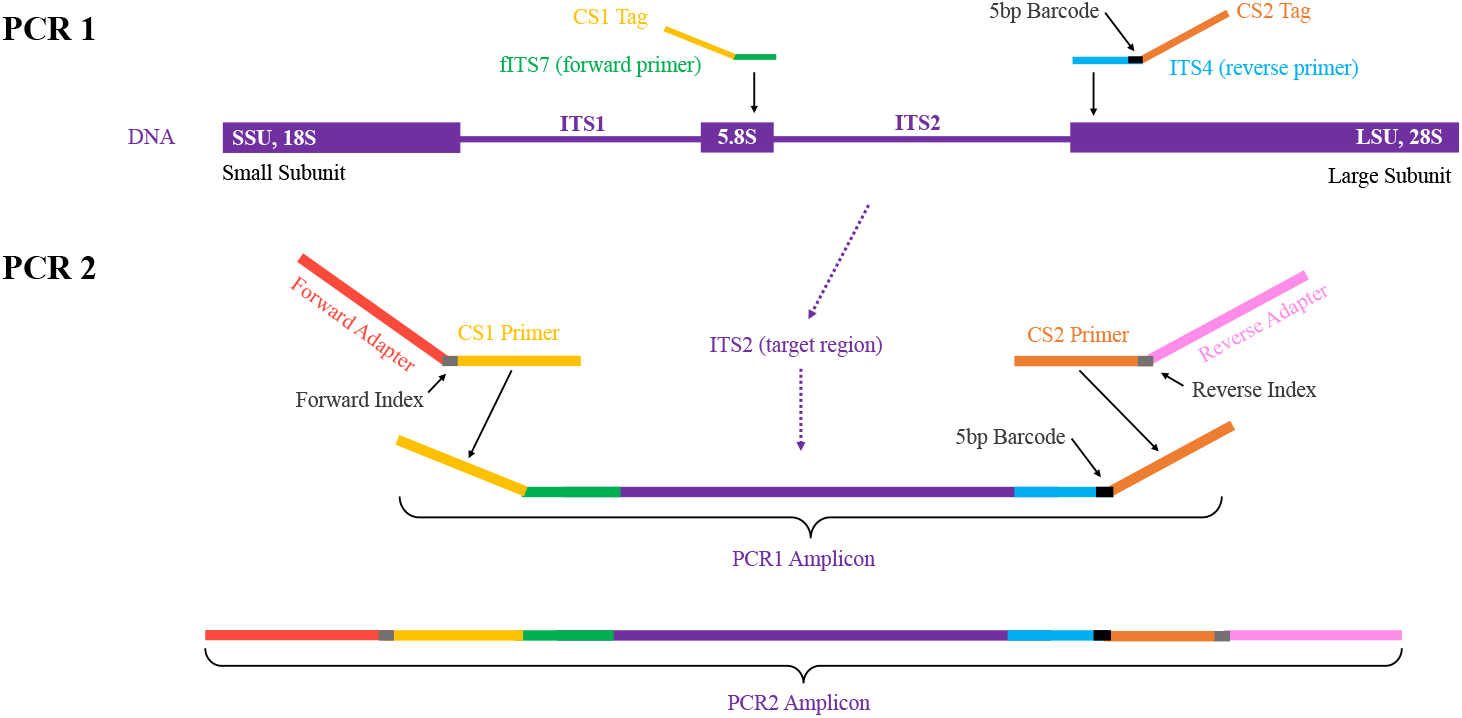
Two-Step PCR for Illumina Sequencing. During PCR1, forward primer fITS7 and reverse primer ITS4 target the nrITS2 region of DNA. These primers include 3’ CS (Fluidigm consensus sequence) tag that provides a priming site for PCR2. Between the ITS4 primer and CS tag is a 5bp barcode (black rectangle). During PCR2, paired-end Illumina adapters prime to the CS tags of the PCR1 product.

## METHODS

### Sampling

We sampled tissues from 766 dried specimens from several fungaria (Table 1, Appendix 1). Most specimens were sampled from the Sam Mitchel Herbarium of Fungi at Denver Botanic Gardens (DBG) and represent macrofungi from the Southern Rocky Mountain region. The specimens sampled represent diverse orders of Dikarya that are summarized in Table 1. A total of 382 specimens belong to the genus *Lactarius* (Russulales). This focused set was chosen for the purpose of evaluating whether the nrITS2 sequence data is capable of recognizing MOTUs at the species rank for a well-sampled genus. Specimens were sampled using forceps that were cleaned with 70% ETOH and KimWipes. A tissue fragment slightly larger than a grain of rice was sampled from a sporocarp’s hymenium and stored in an Eppendorf tube labeled with the specimen catalog number. Specimen information was recorded on an Excel spreadsheet and tracked through DNA preparation and sequencing.

**Table 1.**
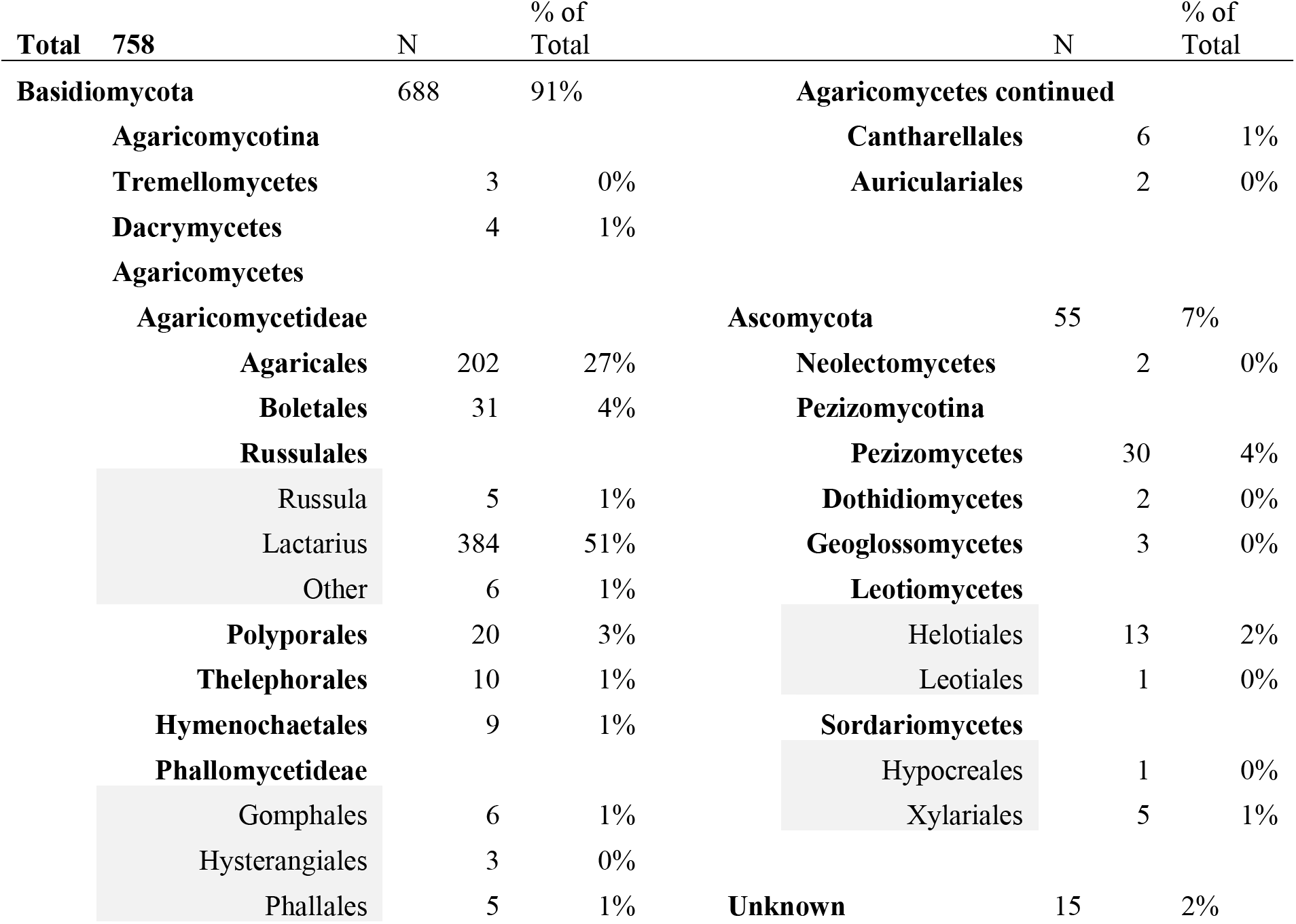
Distribution of Taxa Represented in 766 Specimens primarily from the Sam Mitchel Herbarium of Fungi at DBG. 766 fungal specimens were sampled within subkingdom Dikarya. Taxa are shown by phylum, class, order, and Russulales genera *Russula* and *Lactarius*.

### DNA Extraction

We used two methods of DNA extraction to compare a simple method designed for efficiency and cost-effectiveness to a method designed for DNA quality. The simple method uses a two-step DNA extraction that involves the following: add 200 μL of Extraction Solution (10 ml of 1 M Tris, 1.86 g KCl, 0.37 g EDTA, 80 ml ultrapure H2O, add 1 M NaOH to achieve a pH = ~ 9.5-10.0, 100 ml with ultrapure H2O) and a pinch of sterile sand to each tissue sample; let stand for 10 minutes then freeze samples all the way through to soften tissues (temperatures <0°C will suffice for freezing); thaw samples then grind tissues until homogenized using a sterilized micropestle (Fisher Sci 12-141-368); incubate at 90°C for 10 minutes before adding 200 μL of 3% BSA solution; vortex and centrifuge briefly (30 seconds) to concentrate denser material at the bottom of the tube. Samples are then considered PCR ready where DNA templates for PCR can be drawn directly from the supernatant. The second DNA extraction method followed the materials and instruction from the Omega Bio-tek E.Z.N.A. Fungal DNA Mini Kit (here forward referred to as mini kit extraction).

### Sanger Sequencing

We applied Sanger sequencing for the purpose of cross comparison between sequencing methods. PCR was done in 25 μL volumes: 1.25 μL at 10 pmol concentration each of forward primer ITS1F (Gardes and Bruns, 1993) and reverse primer ITS4 (White et al., 1990); 12.5μL of MyTaq Mix (Bioline: including taq polymerase, PCR buffer, and free nucleotides); 9 μL of PCR pure water; and 1μL of DNA template. PCR thermal cycler steps involve initial denaturation for two minutes at 95 °C, 33 cycles of: denaturation for 30 seconds at 95 °C; annealing at 30 seconds at 55 °C; and extension for two minutes at 72 °C, followed by a final extension at 72°C, the PCR is stopped and chilled at 4°C indefinitely. All PCR products were inspected through 1% agarose gel electrophoresis. Products with no visible bands were not processed further. PCR product producing strong bands were cleaned with a simple ethanol precipitation: 1 vol 100% ETOH; vortex to mix then let stand for 10 min; centrifuge at maximum speed for 2 min; remove supernatant without disturbing pellet; add 200 μL of 75% ETOH; vortex to mix; centrifuge at max speed for 30 seconds; remove supernatant; place tubes on a 65°C heat block to evaporate residual ETOH; add 20 μL of PCR pure water to suspend the DNA. Cleaned PCR product was quantified using a Nanodrop (Thermo Scientific). A minimum of 25 ng of DNA was sent to ELIM Biopharmaceuticals, Inc. (Hayward, CA) for Sanger sequencing using ITS1F and ITS4 primers.

### High-throughput Sequencing

We modified an Illumina MiSeq approach developed for metabarcoding of environmental fungal communities (Ihrmark et al., 2012) in order to sequence numerous macrofungal specimens. Our method can be described as a highly multiplexed approach where barcoded primers are later nested between Illumina indexes in a two-step PCR process outlined in Figure 2. The first PCR step (PCR1) targets the ITS2 portion of the nrITS region using primers fITS7 (Ihrmark et al., 2012) and ITS4 (see above). Attached to the 5’ end of the primers is a variable spacer region followed by a 3’ CS tag (Fluidigm consensus sequence) that provides a priming site for secondary PCR. The variable spacers consist of an additional 0-6 base pairs in the forward primer and 0-4 base pairs in the reverse primer. The modification we introduce to PCR1 is the insertion of a 5 bp barcode between the ITS4 region and the spacer of the reverse primer (black bar in Figure 2). The eight unique 5 bp barcode sequences we introduce are sourced from Shokralla et al (2014). Each DNA sample is assigned one of the eight barcoded primers. PCR1 products – each representing a sample with a unique PCR1 barcode – are pooled to produce a mock community that will be amplified in PCR2 (for PCR details, see below). The PCR2 step uses primers with CS tags that anneal to the complementary sequence of the PCR1 product. Adjacent to the CS tags are unique paired-end Illumina indexes for further multiplexing of samples, lastly followed by Illumina adapters. We performed PCR2 reactions on 94 mock communities of 8 specimens, and two on mock communities of 7 specimens for a total of 766 specimens across 96 reactions. Primer sequence information and instruction on assembling PCR products is provided in Appendices 2 and 3.

PCR1 and 2 use the same recipe as the 25μL reactions described for Sanger. Thermalcycler settings are as follows: PCR1 settings – initial denaturation for two minutes at 95°C, 30 cycles of: denaturation for 60 seconds at 95°C; annealing for 60 seconds at 54°C; and extension for 60 seconds at 72°C, followed by a final extension at 72°C for ten minutes; PCR2 settings −initial denaturation for one minute at 95°C, 15 cycles of: denaturation for 30 seconds at 95°C; annealing for 30 seconds at 60°C; and extension for 60 seconds at 68°C, followed by a final extension at 68°C for five minutes.

We evaluated PCR1 and 2 amplification using 1% agarose gel electrophoresis. PCR1 products were assessed as being in one of three categories – strong, weak, or no amplification. All PCR1 products were cleaned using the same ethanol precipitation method described above but modified for 0.2mL wells in 96-well plate format. PCR1 barcoded products were multiplexed into pools of eight unique barcoded samples. Each PCR product was sampled in 3μL, 5μL, or 10μL volumes based on gel electrophoresis assessments of strong, weak, and non-reactions respectively. This was done to streamline the normalizing of hundreds of PCR products within a pooled mock community or sequencing library. Detailed protocols for all labwork are provided online through https://coloradomycoflora.org/process/protocols/. PCR2 products were then pooled into a single library and sent the University of Idaho’s IIDS Genomics and Bioinformatics Resources Core (GBRC). There the library undergoes fragment analysis, qPCR, and bead based size selection and library amplification before sequencing using Illumina MiSeq V2 (Nano 500 cycles – 250 bp forward, 250 reverse pair ended).

### Sanger Sequence Data

Sequence data received as .abi files was read and edited using default parameters for Geneious v. 2020.2.2. All sequences were trimmed from the ends, removing sequence data with poor electropherogram signal. Contigs were assembled if forward and reverse reads were available for a specimen, except where specimens were sequenced only in the reverse direction (ITS4) to target the ITS2 region. BLAST queries through the NCBI GenBank nucleotide database (nt) were used to assess sequence taxonomy.

### Illumina Sequence Data

Illumina MiSeq V2 (Nano) sequence data were received as demultiplexed fastq.gz files. The GBRC sequencing facility will fully sort reads by PCR1 barcode and PCR2 indexes, placing all sequence reads pertaining to an individual specimen into its own fastq.gz file. We selected the option for Illumina 5’ indexes, adapters, and primers to be removed from sequences by the GBRC.

We evaluated sequence data further using Python scripts to perform a number of bioinformatic tasks (Figure 3). The number of read pairs obtained per specimen was used as one metric for success of a specimen’s sequencing. This was evaluated both before and after quality control (QC) read processing. For each specimen’s read file, any overhanging primers or adapters were removed from the 3’ ends of reads (those with short inserts causing read through), and sequences shorter than 200bp in length were removed, using CutAdapt (Martin, 2011). For the rest of read processing, we then used the DADA2 pipeline (Callahan et al., 2016), a complete Illumina processing tools designed for ecological metabarcode sequencing projects. Following the software’s recommended workflow, DADA2 was run with an initial screen removing low quality reads from the dataset (QC processing, maximum expected error per read parameter 3). We examined the number of read pairs per specimen both from raw data files before processing and also after QC processing with DADA2. We did this to determine if there was a correlation between a successful PCR1 amplification and sequencing as revealed by the number of read pairs (before and after processing) and the Phred scores from a specimen’s read pairs. Then DADA2 was used to model a sequencing error distribution from the data and to apply that model to correct likely erroneous reads. Corrected forward and reverse reads were then merged where possible to produce full PCR fragment sequences (e.g., contiguous sequences or contigs). Bimeras (2-part chimeras) were removed based on fragment distributions using the per-sample option. The final product of this pipeline is a set of unique sequences that have passed these various quality filter steps each with an associated read count reflecting that sequence’s abundance in a singe specimen’s dataset (the specimen’s inference table).

**Figure 3.**
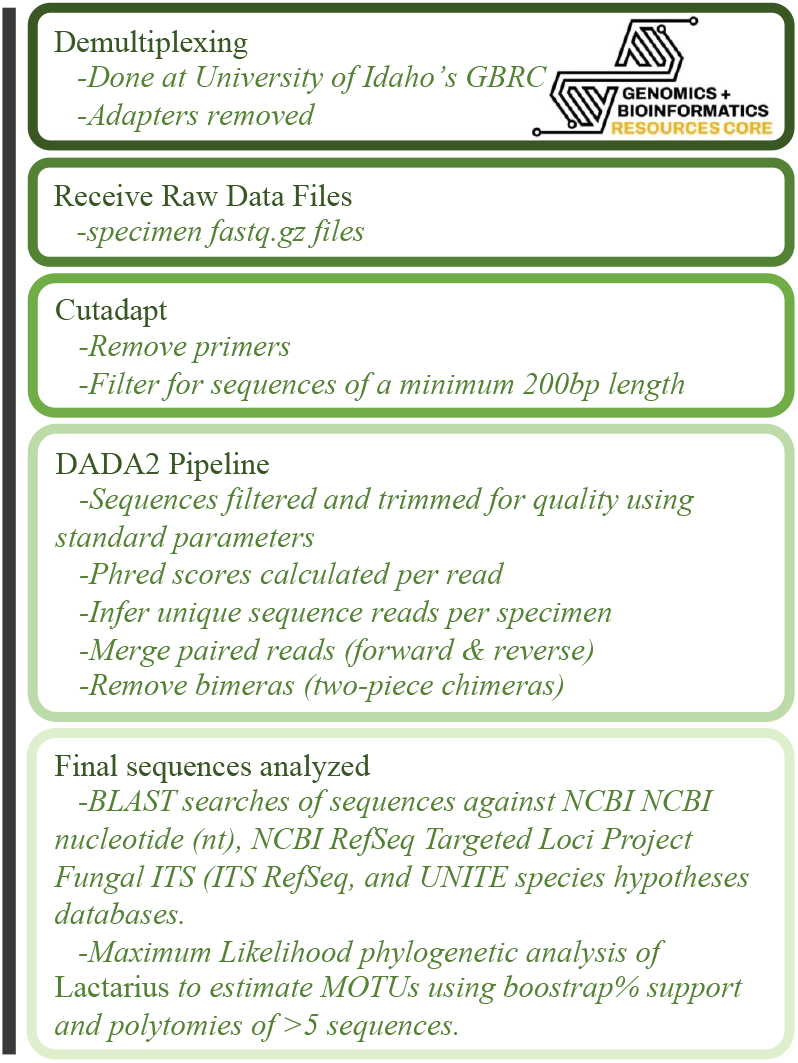
Bioinformatic Pipeline for Processing Illumina Sequence Reads.

The distribution of unique nrITS2 contigs was evaluated for each specimen. More than one unique sequence was present in the processed Illumina data for most specimens, but generally one sequence, consistent with the specimen’s barcode sequence, was much more abundant than the rest in the pool of PCR products. The sequence supported by the greatest number of reads for specimen was chosen as the representative nrITS2 sequence for downstream analysis. The other sequences detected in this specimen’s Illumina data pool include intragenomic variants, contaminants, and potential erroneous sequences that slipped through processing; many of these are at very low abundance in the specimen read pools.

### Sequence Evaluations

Representative sequences from Illumina and Sanger methods were evaluated using BLAST searches of the NCBI nucleotide (nt), NCBI RefSeq Targeted Loci Project Fungal ITS (ITS RefSeq)(Schoch et al., 2014), and UNITE SH (Kõljalg et al., 2005) databases. The proportion of sequences that fell within categories of 97.5-100%, 95-97.5%, 90-95%, 85-90%, and <85% sequence similarity to the top hit in each database was compared to assess the existence of similar sequences to current databases, for both the inclusive but unreliable NCBI nt collection and the well-curated but sparsely populated ITS RefSeq collection. The broader taxonomic rankings across all sequences were evaluated using the UNITE species hypothesis matching analysis tool (github.com/TU-NHM/sh_matching_pub) as implemented on the free online PlutoF data management platform (plutof.ut.ee, data downloaded 10 June 2021) using the ITS2 option and otherwise default settings. Sanger and Illumina sequence data from the same specimen were compared against each other.

### Phylogenetic Analyses

A DNA matrix of *Lactarius* sequence data from Sanger and Illumina sequencing methods. Sequence data from top BLAST hits of the nt, ITS RefSeq, and UNITE databases were also added to the dataset. Because this dataset represented Rocky Mountain species of *Lactarius* we included at least one representative ITS sequence from each species identified in the study by Barge and Cripps (2016). Initial auto-alignment was performed using the MUSCLE v.3.8.31 alignment tool (Edgar, 2004). Additional manual adjustments were performed using JalView (Waterhouse et al., 2009). Phylogenetic analysis was performed in RAxML (Stamatakis, 2014) using the default settings and implemented through the CIPRES Science Gateway (Miller et al., 2010). Rapid bootstrapping analysis was performed using 100 bootstrap replicates. Trees were visualized using FigTree v. 1.4.4 (Rambaut and Drummond, 2019). The phylogeny was used to recognize MOTUs, which were defined as clades with more than three sequences, clades representing specimens with the same species identification, and bootstrap support greater than 75% or the highest supported polytomy with more than five sequences.

## RESULTS

For Sanger sequencing, 337 specimens, including 41 *Lactarius* specimens, used the mini kit extraction method for DNA extraction. Of the total PCR reactions, 274 amplified sufficiently for sequencing (42 *Lactarius*). Of the sequences that came back, 54 were of too low a quality (e.g., bad electrophoresis signal), or were too short after trimming and were therefore discarded. This left 220 sequences (80%) remaining for analysis.

For DNA extraction with Illumina Sequencing, 474 specimens used the simple extraction method and 292 specimens used the mini kit extraction. Evaluation of PCR1 gel electrophoresis showed no discernable difference in amplification success related to extraction method. Evaluation of PCR success showed 136 specimens did not amplify, 203 amplified weakly, and 427 amplified strongly. As expected, the age of the specimen was the most important predictor of PCR1 success where the proportion of specimens that “failed” PCR amplification was greater in older specimens overall (Figure 4). Regardless of PCR success, all 766 specimens were processed for PCR2. All 96 pools of PCR2 reactions produced amplification but because nearly every reaction was a DNA template pool of seven to eight PCR1 samples amplification appeared like faint multiple bands or smears on the gel.

**Figure 4.**
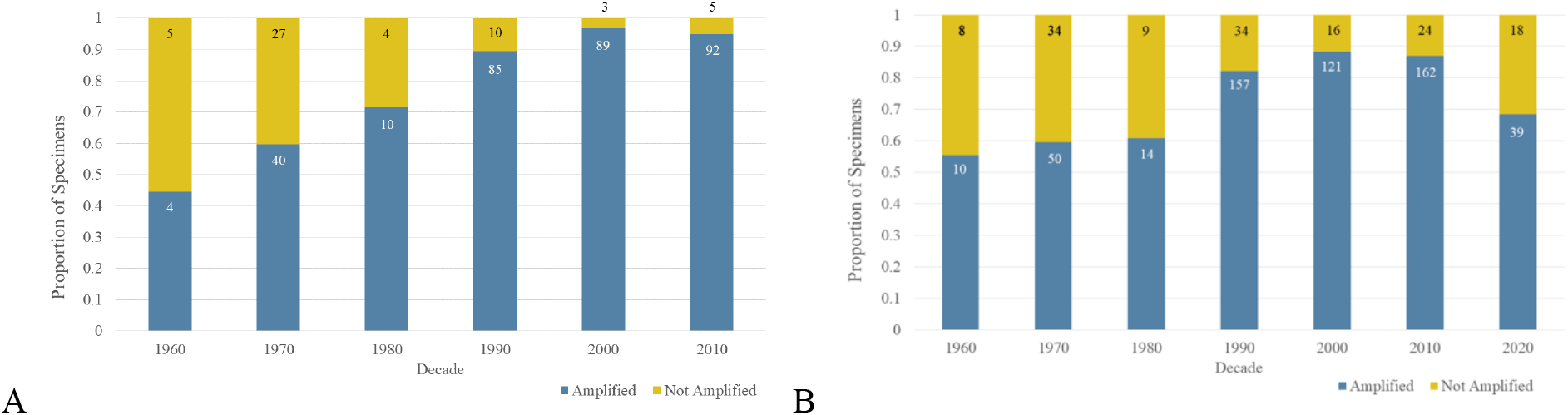
Results of PCR 1 nrITS2 amplification for A. 382 *Lactarius* and B. all 766 macrofungal specimens by decade. Blue represents amplified specimens and yellow represents the non-amplified specimens as a proportion of the total. Samples sizes are annotated as numbers in the amplified/non-amplified portions of each column. Specimens lacking date information were excluded.

When assessing the relationship between PCR success and sequencing success from the raw data, specimens that did not amplify had fewer reads than specimens that did (Figure 5a). Despite this, specimens producing no bands from PCR1 still had an average of 666 forward reads and there was significant overlap in whisker plots representing 95% of the distribution. The result after QC from DADA2 filtering and removal of short reads showed amplified and non-amplified specimens still maintained similar relative read proportions (Figure 5b). Even though the total number of quality reads was reduced by greater than 50%, the means were 355 and 262 reads for positive and negative PCR amplification respectively. These means fall within each other’s respective quartiles, and the whiskers for each plot overlap nearly 100%. Phred score mean (37.44) and median (38) at the end of QC data processing was nearly identical for both the amplified and non-amplified specimens (Figure 6).

**Figure 5.**
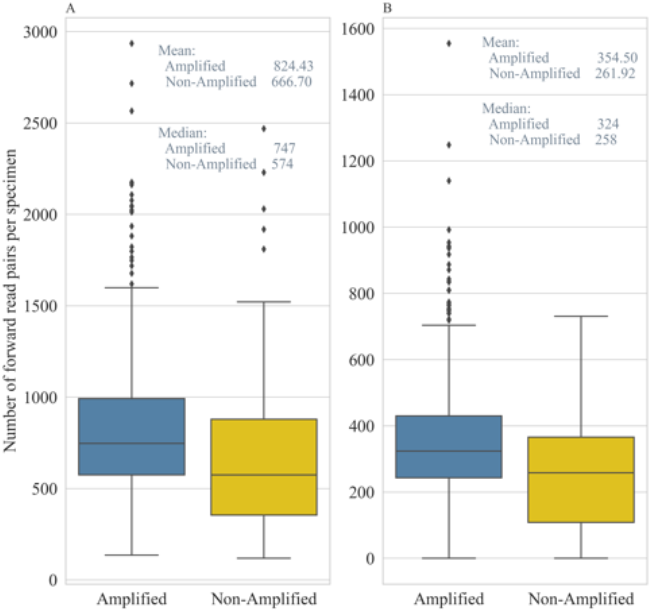
Distribution of forward reads per specimen relative to PCR amplification success. A. Number of reads from raw data. B. Number of reads with DADA2 quality filtering (QC) of data. Specimens that did amplify with strong or weak bands are in blue, and those that did not amplify are in yellow.

**Figure 6.**
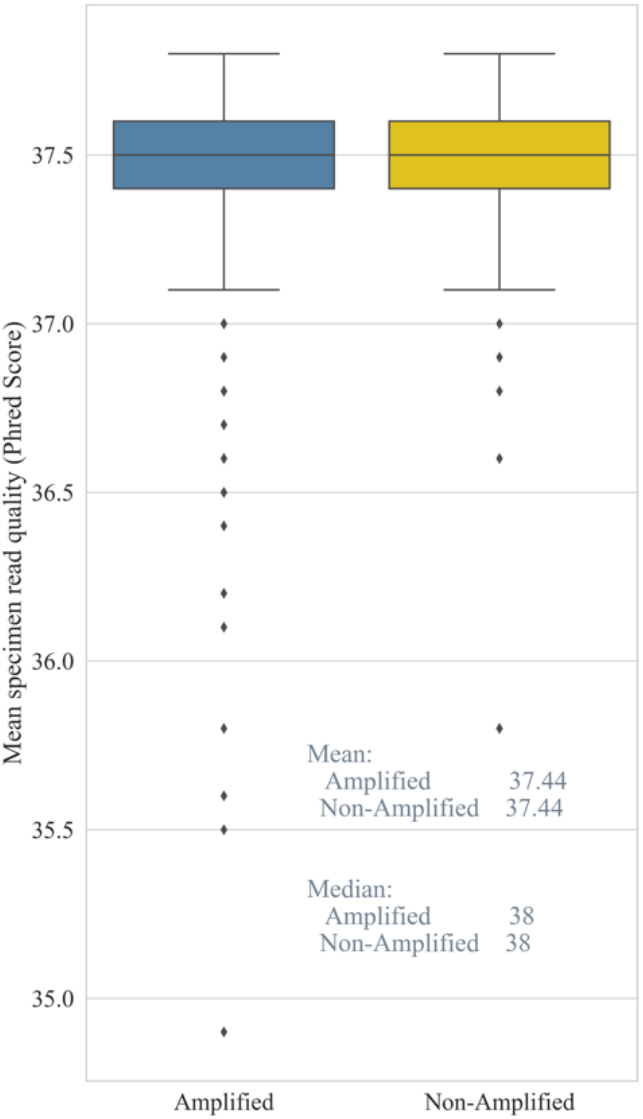
Average Phred scores across all specimens relative to PCR amplification success. Specimens that did amplify with strong or weak bands are in blue, and those that did not amplify are in yellow.

The mean number of read pairs per specimen was 790.2 in the raw data, but dropped to 334.4 read pairs per specimen after QC was completed (Figure 7). Before removal of poor-quality forward and reverse reads from the raw data files, mean Phred score was ≤36.5 (Figure 8). In dropping poor quality reads, the mean forward and reverse read Phred score increased nominally to 37.44 and 37.26 respectively.

**Figure 7.**
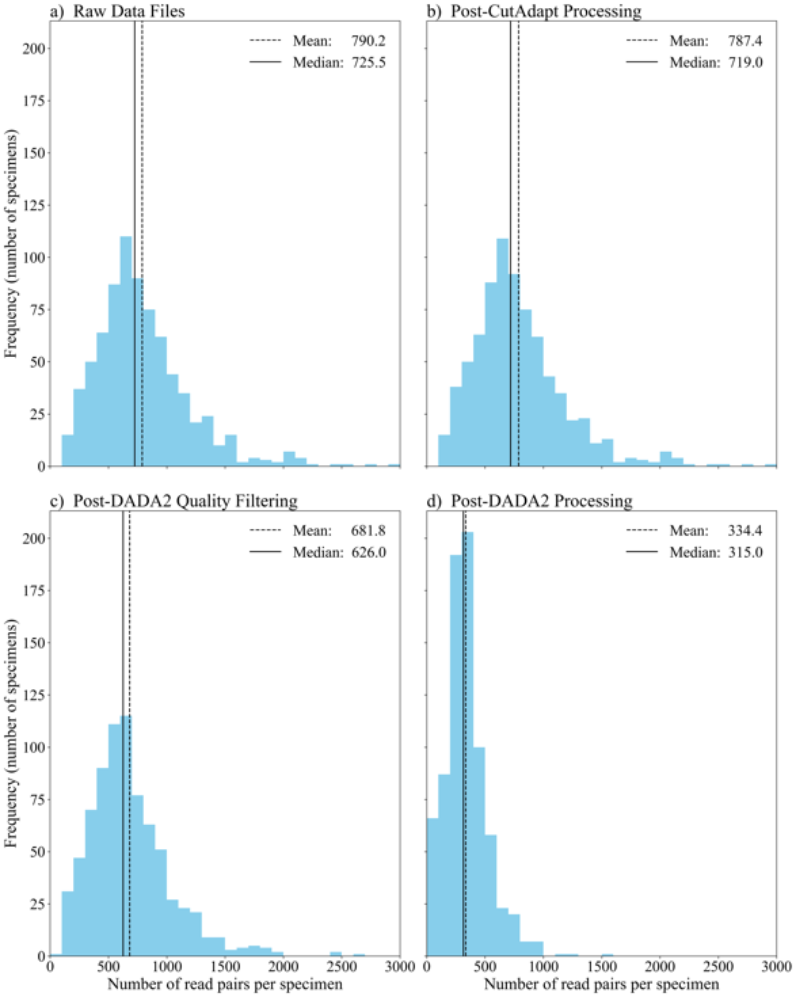
Distribution of Read Pairs per Specimen for 766 Specimens at Four Stages of Processing. Starting with the raw sequence reads received from iBest, three benchmarks during data processing (after processing with both CutAdapt and DADA2) show the decrease in average and median number of read pairs per specimen as lower quality reads are trimmed and/or cut out of the dataset.

**Figure 8.**
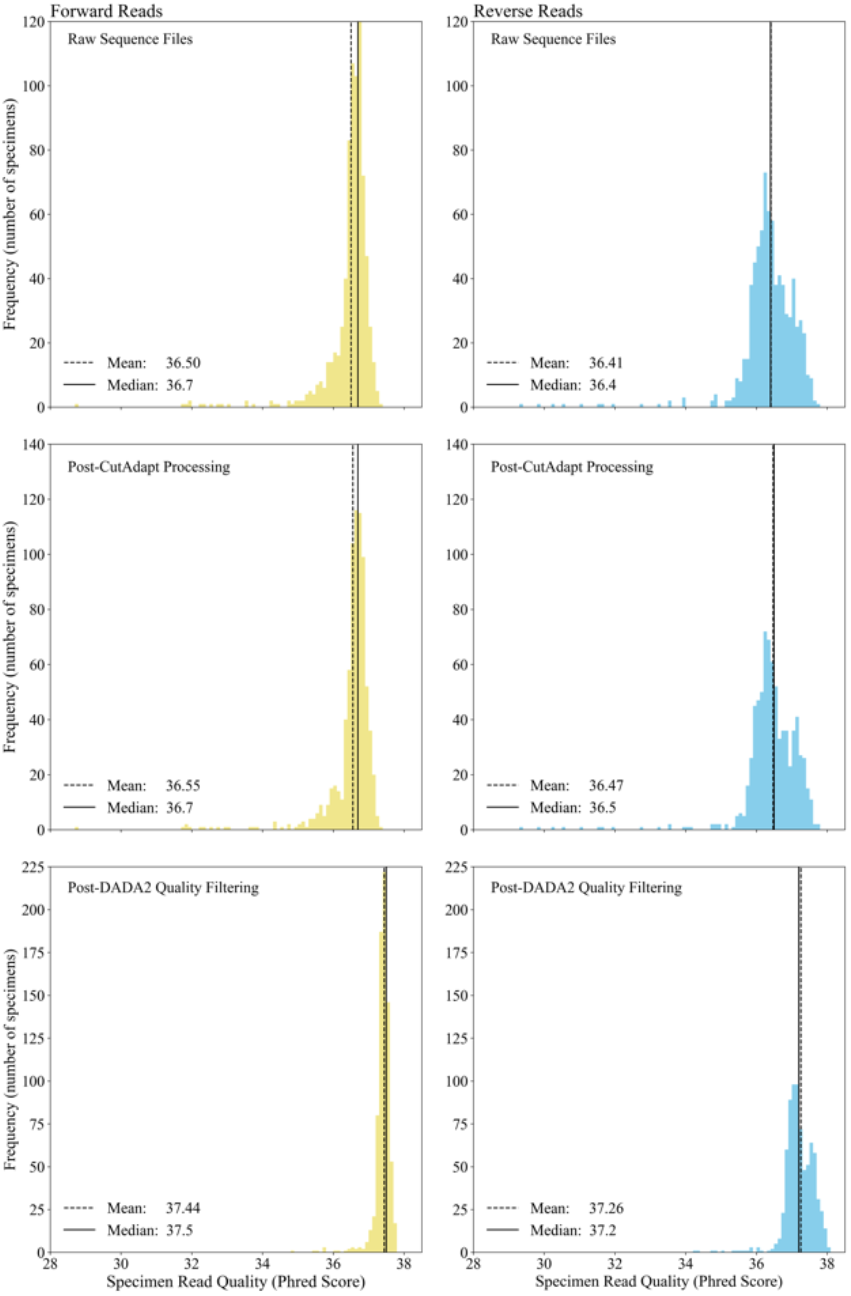
Distribution of average Phred scores for 766 specimens. The average quality scores were from a) raw data files (a), after CutAdapt processing, and going through the last stage in the process after DADA2 quality filtering. Distributions of all forward reads are in yellow, and all reverse reads are in blue.

Four of the 766 specimens failed QC. This represents a failure rate of 0.5%. Three of these four specimens had not amplified during PCR1. In other words, of the total 136 specimens that did not amplify, 133 still produced usable sequence data. When comparing data from Sanger and Illumina methods, Sanger produced 220/274 useable sequences (80%), while the modified metabarcoding method produced 762/766 sequences (99.5%)(Figure 9). In total, sequence data from 762 specimens had read pools that not only passed QC but were large enough to select a representative sequence after contigs of the forward and reverse reads were created. The most abundant sequence in a specimen’s read pool had a mean of 214.4, with a median of 201 (Figure 10).

**Figure 9.**
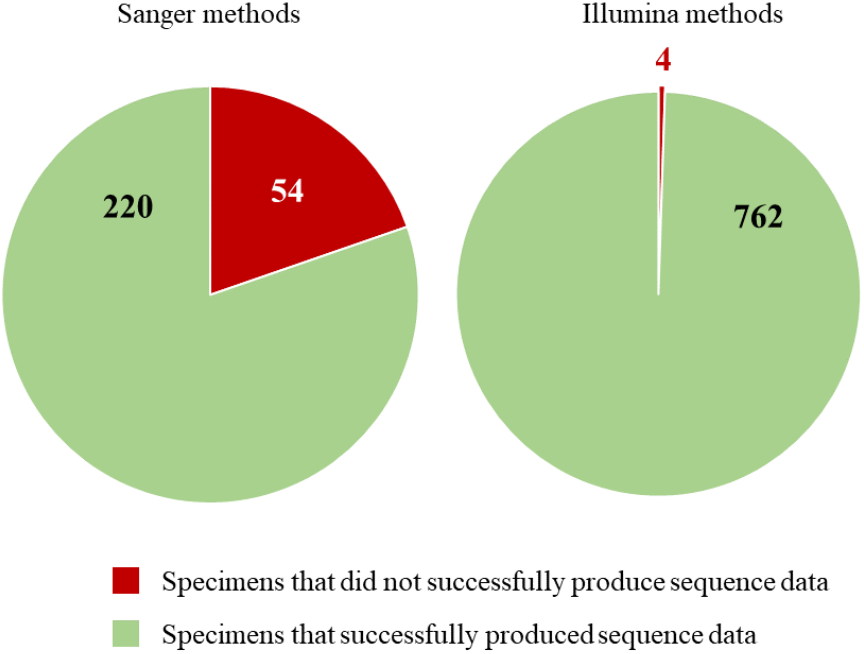
Proportion of specimens between Sanger (A) and Illumina Sequencing methods (B) that were successful (green) in producing sequence data, or unsuccessful (red) in that sequence data was noisy or otherwise low quality.

**Figure 10.**
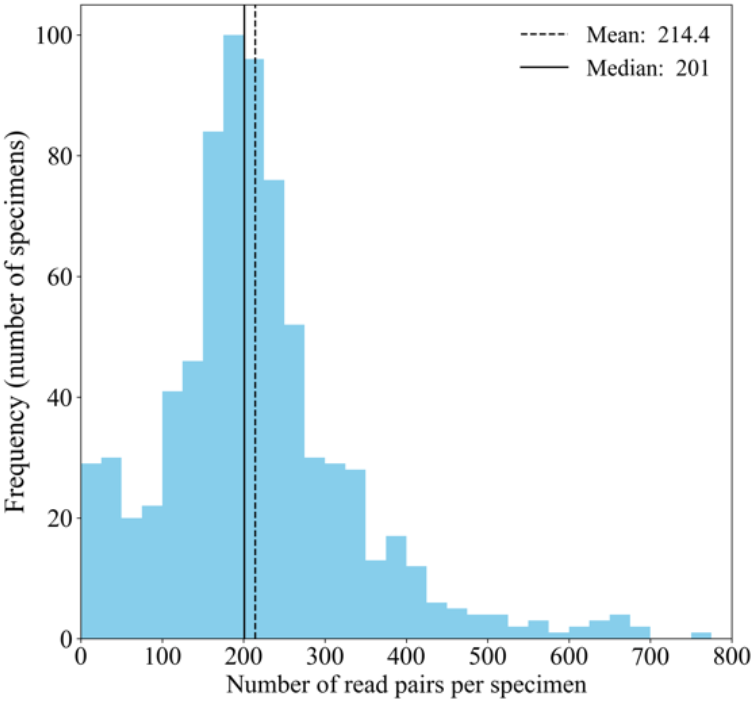
Distribution of the number of contigs (read pairs) representing most abundant recovered from each specimen for all 762 specimens.

To test if Sanger-sequence data matched Illumina data from the same sequences, BLAST alignments were created and compared for % identity and % coverage. The results demonstrated at or near 100% identity between the majority of sequences in Sanger-Illumina BLAST alignments (Figure 11). In addition, over 80% coverage was observed between these sequence alignments.

**Figure 11.**
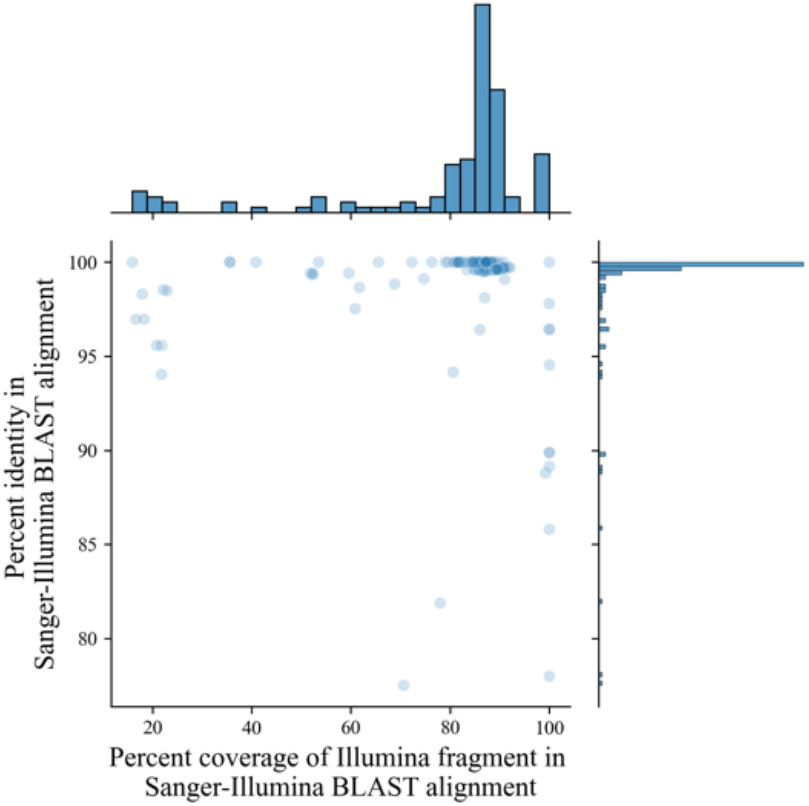
Percent identity match and overlap of Sanger-Illumina sequence BLAST alignment. All Sanger-sequenced specimens were aligned with their Illumina sequence using BLAST alignment. The vast majority of specimens aligned at near-100% identity match, and there was mostly over 80% overlap of the Illumina sequence on the alignment sequence.

Figure 12 shows the distribution of data as presented in the PlutoF data management platform (plutof.ut.ee). By comparing the taxonomic representation of specimens sampled for this study (Table 1) 50% of sequences match the genus *Lactarius*, with *Amanita* and *Laccaria* as the second and third most abundant genera represented. 87% of specimens belong to phylum Basidiomycota, and 8% to Ascomycota. These proportions from the original specimen data are supported by and match the species hypothesis BLAST search results. Results of percent similarity and percent sequence overlap to the top match from each of these databases is provided in Appendix 4.

**Figure 12.**
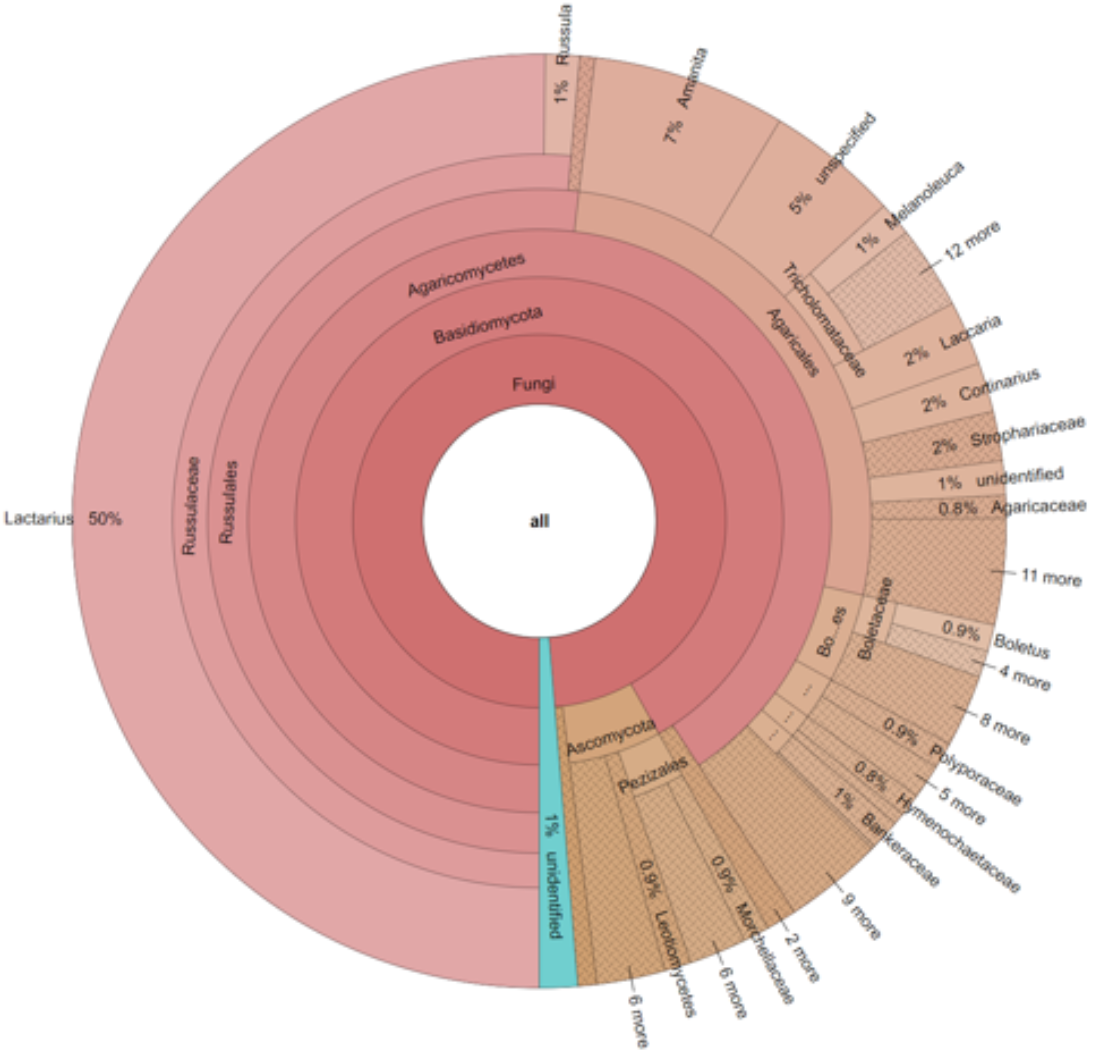
UNITE species hypotheses (SH) found in 762 ITS2 Illumina sequences. Matches produced using a development version analysis in PlutoF data management platform (pluto.ut.ee).

We used Genbank and UNITE to investigate species hypotheses and to explore the taxonomic identity for sequences generated in this study. The sequence data produced from macrofungal specimens had varying representations across databases with the NCBI nt database having the highest scoring and the ITS RefSeq database the lowest scoring hits (Figure 13). Upon closer evaluation of taxonomic identities returned from searches of reference databases, 87 of the 766 results were not consistent with the taxonomic IDs of the specimens. Of these, 28 can be attributed to specimen misidentification or nomenclature changes not reflected in the specimen or sequence record. This leaves 59 sequences (8% of all specimens) that do not reliably match the specimen’s current identity. Whether this is due to contamination or from sampling the wrong unique sequence from a specimen’s read pool remains to be investigated. Accession numbers for all sequences uploaded to Genbank are provided in Appendix 1 (FOR REVIEW sequences are provided in fasta format in Appendix X).

**Figure 13:**
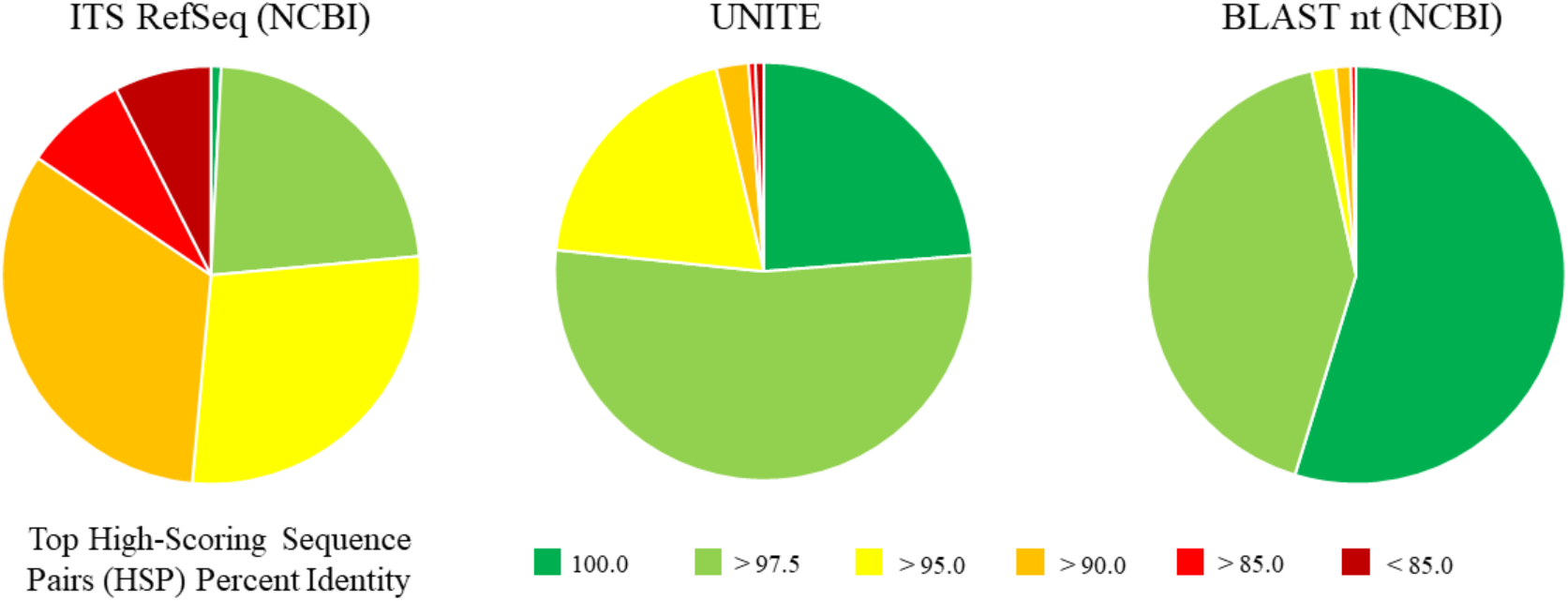
Results of sequence matches to major sequence databases.

Phylogenetic analysis of *Lactarius* ITS2 sequence data generated in this study used a data matrix of 487 sequences. In total, 377 sequences were produced using Illumina MiSeq, 28 were produced using Sanger, and 82 were sourced through GenBank (Appendix 1). The phylogeny resulted in approximately 20 *Lactarius* MOTUs from the Rocky Mountain region (Figure 14). Of these, 17 have MS bootstrap support ≥75%. Two of these have 100% bootstrap support, five have 95-99%, seven in the 85-94% range, and two between 75-84%. Three are below 75% but above 55%. These are labeled as *L*. aff. *hepaticus, L*. aff. *alpinus*, and *L. scrobiculatus* in the phylogeny. The failure to resolve these taxa may be due to incomplete representation of taxa within each clade, or lack of differentiation within the nrITS2 region in this region of the phylogeny. Further sequence data from other molecular markers would be the next step in identifying what limits this dataset from providing better resolution for MOTUs.

**Figure 14.**
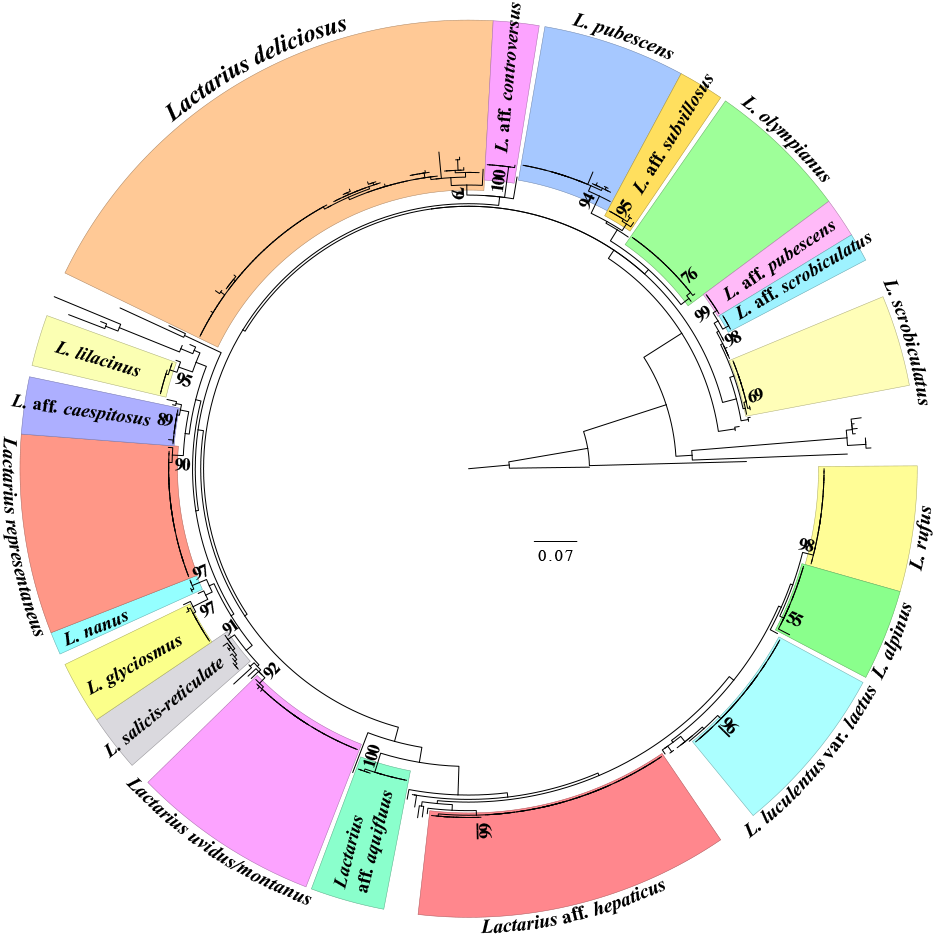
Maximum Likelihood RAxML-NG phylogeny of 487 *Lactarius* ITS2 sequences. Newly generated sequences consist of 378 from Illumina MiSeq Nano and 28 from Sanger sequencing methods. The remaining 82 sequences represent data from GenBank representing Rocky Mountain *Lactarius* taxa. A total of 20 clades are represented by > 3 sequences and are identified as MOTUs. This is based on resolution, bootstrap support, and the proportion sequences from named specimens within a clade. Numbers next to branches indicate ML bootstrap percentages from 100 replicates.

## DISCUSSION

This project uses a nested barcode primer in a two-step PCR approach for metabarcoding fungal communities to produce a method for the mass sequencing of macrofungal specimens. This approach deals with issues of scale, effectiveness, and cost that have largely limited large-scale DNA barcoding of fungarium specimens. The ability to sequence more specimens will provide researchers with the opportunity to more effectively study fungarium collections for systematic and taxonomic research.

The DNA extraction methods tested in this study did not differ in their ability to provide sequence data. This supports the cost-effective simple DNA extraction method for processing 100’s of specimens. Costs can be further leveraged when using the Illumina MiSeq Nano. The MiSeq Nano’s flat rate for sequencing means that the cost per nrITS2 sequence goes down as you scale up the number of specimens (Figure 1). The one caveat is that the MiSeq Nano kit we used is limited to 800,000 cycles. This would limit the number of specimens you can scale up before the number of reads per specimen begins to affect sequence reliability. As a result, we suggest limiting a MiSeq Nano sequencing run to 800-850 specimens. If sequencing of more specimens (>1000) is desired, then a MiSeq V3 kit with 10 million cycles is recommended.

The high-throughput sequence data demonstrated several advantages over Sanger. One example was the ability to produce nrITS2 sequence data from specimens that did not amplify during PCR (Figure 5). Of the specimens sent for sequencing 99% produced nrITS2 sequence data that passed QC. In contrast, the specimens that did not amplify after PCR were not sent out for Sanger sequencing. Of the ones sent for Sanger sequencing, only 80% produced usable sequence data (Figure 9). When comparing Illumina sequence to Sanger sequence data, there was a near 100% match in sequence identity (Figure 11). One caveat to this evaluation is that Illumina sequence data targeted nrITS2, whereas Sanger can target the whole nrITS region.

More evaluation of the data is needed, but the specimens that were sampled to represent diversity within the Dikarya were recovered proportionally (compare Table 1 and Figure 12). In evaluating age of collections, we were able to recover sequence data from specimens collected between 1910-2020. This suggests this method has broad application for fungaria specimens representing taxonomic and temporal diversity. The potential of this application on other collections (e.g., culture collections, herbarium specimens, etc.) is promising, but remains to be explored.

Phylogenetic analysis of 487 *Lactarius* ITS2 sequences was considered effective in producing MOTUs. Figure 14 has a total of 20 highlighted clades represented by 4 or more sequences. Seventeen of these have maximum likelihood bootstrap percentages >70%. Of the three with bootstraps <70%, only *L. scrobiculatus* formed a clade with sequence data from Genbank. *Lactarius* aff. *alpinus* (MLBS 55%) and *L*. aff. *hepaticus* formed clades that were made of entirely original sequence data from DBG specimens. While these clades are not well supported, these and the other clades of *Lactarius* from the Southern Rocky Mountain region will be the subject of further systematic and taxonomic study using morphology and additional sequence data.

The nrITS region is considered a valuable tool in generating MOTUs in fungi when used to produce species hypotheses (Kõljalg et al., 2013; Osmundson et al., 2013). Regardless, it is important to understand the limits in the nrITS to resolve taxonomic and systematic problems (Lindner and Banik, 2011; Hilário et al., 2021). These must be understood especially when only using a portion of the region to study diversity questions in fungi.

The flood of fungal sequence data from environmental metabarcoding studies has expanded our understanding of fungi in the environment, it has also highlighted the number of unknown fungal taxa (e.g., “Dark Taxa”)(Tedersoo et al., 2017; Ryberg and Nilsson, 2018). This problem is exacerbated by the fact that over 70% of fungal species do not have a representative sequence available on fungal sequence databases (Hawksworth and Lücking, 2017). Sequence data from fungarium specimens can resolve these issues of undescribed environmental diversity by enhancing sequence representation from vouchered specimens in sequence databases. The potential for fungarium specimens to identify unknown fungi from eDNA samples has been demonstrated in a study of *Mortierella* (Nagy et al., 2011).

The ability to sequence all the specimens of a single taxon within a collection has the benefit of resolving misidentified specimens and identifying cryptic species. The study by O’Donnell et al. (2011) on the genus *Morchella* demonstrates the significance of cryptic species in macrofungi. Similar studies have also shown cryptic diversity in North American *Cantharellus* (Buyck and Hofstetter, 2011; Foltz et al., 2013; Leacock et al., 2016). The ability to generate sequence data more broadly in fungal collections will help advance the use of fungaria in filling in the gaps of fungal diversity.

Lastly, this approach provides a new opportunity to address the DNA barcode gap between many fungal species. When evaluating the nrITS between two species it is necessary to produce sufficient data to understand both inter and intra-species variation. The ability to sequence all the specimens of a taxon provides an opportunity to understand the abilities of the ITS2 as a proxy for the fungal DNA barcode.

## ACKNOWLEDGEMENTS

We wish to thank Dr. Bryn Dentinger at the Natural History Museum of Utah and the University of Utah for providing the discussion and ideas that inspired this research. We appreciate Dan New and Matt Fagan at the University of Idaho for initial consulting and additional library conditioning and sequencing. Data collection and analyses performed by the IIDS Genomics and Bioinformatics Resources Core at the University of Idaho were supported in part by NIH COBRE grant P30GM103324. AWW wishes to thank Dr. Peter Kennedy for initial advice and guidance on the metabarcoding methods in fungi. We wish to express appreciation to the Daniel E. Stuntz Memorial Foundation (stuntzfoundation.org) for the Mycology Grat awarded to JJL that provided funding for sequencing and analysis of Idaho specimens. Phylogenetic analyses were implemented through the CIPRES Science Gateway (phylo.org) which is supported by NIH award 5R01GM126463-01, with day to day operations supported by NSF DBI award 1759844.

